# Dose range-finding toxicity study in rats with recombinant human lactoferrin produced in *Komagataella phaffii*

**DOI:** 10.1101/2024.01.22.576753

**Authors:** Ross Peterson, Robert B. Crawford, Lance K. Blevins, Norbert E. Kaminski, June S. Sass, Bryce Ferraro, Roma Vishwanath-Deutsch, Anthony J. Clark, Carrie-Anne Malinczak

**Author notes:** Ross Peterson, PhD: Director of Regulatory Affairs, Helaina Inc. New York, NY, USA; ORCID: 0009-0002-7405-4602. Robert B. Crawford: Institute for Integrative Toxicology, Michigan State University, East Lansing, MI, USA. Lance K. Blevins, PhD: Assistant Professor, Institute for Integrative Toxicology, Michigan State University, East Lansing, MI, USA; ORCID: 0000-0002-8911-8843. Norbert E. Kaminski, PhD: Professor, Department of Pharmacology & Toxicology, Director and Food and Consumer Product Ingredient Safety Endowed Chair, Center for Research on Ingredient Safety, Director, Institute of Integrative Toxicology, Michigan State University, East Lansing, MI, USA; ORCID: 0000-0002-2144-428X. June S. Sass, BS: Research Associate, Nutritional Biology & Safety, Helaina Inc. New York, NY, USA; ORCID: 0009-0002-6397-4614. Bryce Ferraro, BS: Lead Engineer, Early-Stage R&D, Helaina Inc. New York, NY, USA; ORCID: 0009-0003-8259-3478. Roma Vishwanath-Deutsch, MS: Scientist, Nutritional Biology & Safety, Helaina Inc. New York, NY, USA; ORCID: 0009-0003-0236-4270. Anthony J. Clark, PhD: Chief Technology Officer, Helaina Inc. New York, NY, USA; ORCID: 0009-0002-9425-0221. Carrie-Anne Malinczak, PhD: Head, Nutritional Biology & Safety, Helaina Inc. New York, NY, USA; ORCID: 0000-0003-3377-5921. **Corresponding author information:** Carrie-Anne Malinczak, PhD.; Address: 345 Park Ave, South, New York, NY 10010, USA.

## Abstract

The oral toxicity of recombinant human lactoferrin (Helaina rhLF, Effera™) produced in *Komagataella phaffii* was investigated in adult Sprague-Dawley rats by once daily oral gavage for 14 consecutive days. The study used groups of 3-6 rats/sex/dose. The vehicle control group received sodium citrate buffer and the test groups received daily doses of 200, 1000 and 2000 mg of rhLF per kg body weight. Bovine LF at 2000 mg/kg body weight per day was used as a comparative control. Clinical observations, body weight, hematology, clinical chemistry, iron parameters, immunophenotyping, and gross examination at necropsy were used as criteria for detecting the effects of treatment in all groups and to inform dose levels for future toxicology studies. Quantitative LF levels were also analyzed as an indication of bioavailability. Overall, administration of Helaina rhLF by once daily oral gavage for 14 days was well tolerated in rats at levels up to 2000 mg/kg/day, or 400x Helaina’s intended commercial use, and indicating that a high dose of 2000 mg/kg/day is appropriate for future definitive toxicology studies.

## INTRODUCTION

Human lactoferrin (hLF), isolated by Bengt Johannsonn in 1960,^1^ is an 80 kD glycoprotein member of the transferrin family of iron binding proteins, made up of a single chain polypeptide of 691 amino acids.^2^ Lactoferrin has received attention due to its bioactive (i.e., functional) properties and beneficial interactions with the immune system.^3^ It is ubiquitous in the human body, present in human biological fluids and mucous secretions (e.g., human milk, tears, and saliva) as well as in the granules of circulating neutrophils.^4^ The concentration of hLF is highest in human milk (hmLF), from 5–7 mg/mL in colostrum to 1–3 mg/mL in mature milk.^5^ While breastfed babies are exposed to high levels of hmLF in early life,^6^ adult humans are also constantly exposed to hLF naturally occurring in tears, saliva and other epithelial cell secretions.^7^ Bovine LF (bLF) is also routinely consumed through ingestion of cow’s milk and was first isolated by Sorensen and Sorensen in 1939,^8^ Since the discovery of LF, studies have been performed in vitro, in experimental animals, and in humans to analyze the multifunctional roles which include, but are not limited to, iron transport and interaction with the immune system.^3,9^

Bovine LF isolated from cow’s milk is generally recognized as safe (GRAS) for various intended uses in conventional food and infant formula, and it is available as a dietary supplement, taken orally for its nutritional benefits.^10^ Literature has demonstrated improved hematological parameters with daily oral administration of bLF in treating iron deficiency anemia in pregnant and non-pregnant women.^11–13^ However, hmLF has been shown to digest more slowly and has stronger affinity for the human LF receptor in the small intestine, indicating that it may be more bioavailable than bLF.^14–16^ Therefore, human forms of LF have the potential to be more functionally active than bLF as a food ingredient. The use of hmLF isolated from human milk has not been pursued due to economic, ethical, and sustainability reasons. Approximately one liter of human milk is required to yield 100 mg of hmLF^17^; not only is this practice costly and wasteful, but milk used for this purpose could be unsuitable for feeding infants. The expression of recombinant proteins provides a method for production at commercial scale with prospective sustainability benefits, and at a lower cost and without ethical concerns. Recombinant human LF (rhLF) has been produced by several biological expression systems including rice,^18^ transgenic cows,^19^ and yeast.^20–22^

Several general toxicity animal studies have evaluated the safety of previously developed rhLF products following oral ingestion. Cerven et al. used a rat model to demonstrate no toxic effects at 1800 mg/kg body weight per day, for 28 days, of rice-derived apo-rhLF (iron unsaturated form).^23^ Another study by the same group found the no observed adverse effect level (NOAEL) of holo-rhLF (iron saturated form) to be greater than 1,000 mg/kg body weight per day.^24^ In a 13-week oral toxicity study in rats, Appel et al. found the NOAEL of rhLF from transgenic cattle to be at least 2,000 mg/kg body weight per day.^25^ Finally, Zhou et al. found that milk powder with rhLF derived from transgenic cattle was as safe as conventional milk powder in a 90-day toxicity study in rats.^26^ These studies provide support for safety regarding the ingestion of rhLF in vivo.

However, few previous studies have examined changes in immune function by rhLF that provide insights into its putative immunotoxicologic effects, crucial for ensuring safety. To our knowledge, no published animal study on rhLF has included comprehensive evaluation of its effects on the frequency of major immune cell types by immunophenotyping. Immunophenotyping is an application of flow cytometry to detect proteins of interest on a single-cell level^27–29^ and it is one of the suggested models to identify compounds of potential immunotoxic risk.^30^ Given that reductions in the frequency of immune cell types can be an indicator of changes to immune function, immunophenotyping is a valuable technique, in combination with other safety endpoints including hematology and clinical chemistry, to characterize the toxicity profile of novel food ingredients and was addressed in the current work.

Helaina Inc. (New York City, New York, USA) has developed rhLF, derived from a glycoengineered yeast, that is substantively similar to hmLF.^31^ Specifically, a proprietary technology involving disruption of an endogenous glycosyltransferase gene (OCH1) and a stepwise introduction of heterologous glycosylation enzymes, enables a modified strain of *K. phaffii* to produce rhLF (Effera™).^32,33^ Despite the structural similarities of rhLF to hmLF, safety of recombinant proteins intended for food use must be demonstrated.

Given that human infants are exposed to naturally occurring hmLF in human milk, and that adults are exposed to hLF in other biological secretions (e.g, saliva), humans are expected to be tolerant to hLF and the addition of Helaina rhLF to the diet should be equally tolerated and not pose a safety risk at appropriately evaluated exposure levels. Tolerability and toxicity of rhLF was evaluated in a 14-day oral gavage dose range-finding study in rats. Endpoints included observable changes in the animals (e.g., behavior, body weight), hematology, clinical chemistry, iron parameters, immunophenotyping, and LF detection in the serum. The goal of the current study was to examine possible toxicological effects of rhLF compared to bLF, which has GRAS designation as a food ingredient, as well as to inform dosage levels for future, longer-term definitive toxicology studies.

## MATERIALS & METHODS

### Test Substances

Recombinant human lactoferrin (rhLF, Helaina Inc.), alpha isoform protein id P02788 of the UniProt database, was produced by fermentation and expressed from a *Komagataella phaffii* recombinant system, purified by microfiltration/diafiltration and cation exchange chromatography, and then spray-dried to a powder. The purity of rhLF was measured by high performance liquid chromatography (HPLC) and sodium dodecyl sulfate–polyacrylamide gel electrophoresis (SDS-PAGE) and is typically greater than 98%. The iron saturation was determined to be 58% saturated.^31^ Recombinant human LF was reconstituted and diluted into sodium citrate buffer (pH 5.5) prior to use.

Bovine lactoferrin (bLF) isolated from bovine milk was purchased from Lactoferrin Co, Australia (Product 11683) and reconstituted and diluted into sodium citrate buffer (pH 5.5) prior to use. Purity, provided by the supplier, was greater than 95% and the iron saturation was 9.9%.

### Animals and Treatment

Nine-week-old CD® (Sprague Dawley) IGS rats were received from Charles River Laboratories, Kingston, New York. Following a seven-day acclimatization period, 42 healthy male and 42 healthy female rats were randomly selected and assigned to treatment and control groups using a computer randomization process. The body weight range prior to the start of dosing was 250-297 g for males and 206-231 g for females.

The animals were socially housed 2-3/cage in solid bottom cages with nonaromatic bedding and appropriate animal enrichment, adhering to the USDA Animal Welfare Act (9 CFR, Parts 1, 2 and 3)^34^ and as described in the Guide for the Care and Use of Laboratory Animals (NRC, eighth edition).^35^ After assignment to study, these animals were implanted with microchips. The individual animal number, implant number, and the Testing Facility study number comprised a unique identification for each animal. The animal cage was identified by the study number, animal number, group number, and sex. Block Lab Diet® Certified Rodent Diet #5CR4, PMI Nutrition International, Inc. was provided ad libitum except for the fasting period on the day of designated procedures. Fresh tap water, which was analyzed routinely for impurities, was available ad libitum. During the time frame of this study, there were no impurities found in the food or water which would affect toxicologic evaluation. The animal room had a 12-h light/dark cycle, only interrupted intermittently for study-related activities. Temperature (68-79℉) and humidity (30-70%) of the room was maintained according to the testing facility SOP.

Recombinant human LF (rhLF, Helaina Inc.), bLF or control vehicle (sodium citrate) was administered orally, once daily by gavage, using a syringe and appropriately sized plastic feeding implement. There were ten dosing groups with 3-6 female and 3-6 male animals in each group. All doses were given at a volume of 10 mL/kg. Three groups of animals (3/sex) received rhLF in sodium citrate at levels of 200, 1000, or 2000 mg/kg bw/day; one group of animals (3/sex) received bLF at 2000 mg/kg bw/day, and one group of animals (3/sex) received sodium citrate vehicle only for the toxicological (main) phase of the study. Three additional groups of animals (6/sex) received rhLF in sodium citrate at levels of 200, 1000, or 2000 mg/kg bw/day, one additional group of animals (6/sex) received bLF at 2000 mg/kg bw/day, and one additional group of animals (3/sex) received sodium citrate vehicle only for the LF absorption phase of the study. The test proteins were mixed daily with sodium citrate (50 mM) buffer, pH 5.5 to the proper concentration and continuously mixed during the dosing procedure. Individual dose volumes were adjusted twice weekly based on body weight.

### Animal Observations

All animals were observed once daily for cage-side observations and twice daily for morbidity and mortality. Detailed clinical observations were performed twice weekly. Body weights were recorded pretest, twice weekly, and on the day prior to study termination.

### Terminal Procedures

Animals were euthanized by carbon dioxide inhalation followed by a Testing Facility SOP approved method to ensure death, (e.g., exsanguination). All animals that underwent scheduled terminal procedures were subjected to a necropsy examination that included evaluation of the carcass and musculoskeletal system, all external surfaces and orifices, cranial cavity and external surfaces of the brain, and thoracic, abdominal, and pelvic cavities with their associated organs and tissues. A veterinary pathologist, or other suitably qualified personnel, was available for consultation during scheduled necropsy. All animals were examined carefully for external abnormalities including palpable masses.

### Clinical Pathology

Blood samples were obtained from each study animal after terminal necropsy via vena cava venipunctures. Clinical chemistry and hematologic parameters are summarized in Supplemental Table 1.

### Lactoferrin Absorption into Serum

Blood (0.3 mL) was collected from each animal via sublingual or other suitable vein at the following time-points on Days 1 and 14: 0.5, 1, 3, 6, 12, and 24 hours from Groups 7-10, and only at hour 1 for Group 6 (vehicle control). Blood was allowed to clot prior to centrifugation to separate serum from the cellular clot. Serum was carefully removed and transferred to a collection tube and stored at ≤-20℃ until ELISA analysis for LF quantification.

#### Human rhLF ELISA

A direct sandwich ELISA method, conducted at Helaina Inc., was used to determine the concentration of rhLF in the rat serum. 96-well plates were coated overnight with capture antibody (100 µg/mL; lactoferrin goat anti-human polyclonal, Bethyl Labs, Cat: A80-143A). The wells were washed and blocked using fish gelatin and incubated for 2 hours. Standards (3.6-210 ng/mL) were prepared in 1:10 rat sera. Plates were washed and standards, positive control (known concentration of rhLF spiked into 1:10 rat sera), samples (diluted 1:10), and blanks (1:10 rat sera) were added to appropriate wells, and plates were incubated for 1.5 hours at room temperature. Plates were washed, a 1:5,000 dilution of detection antibody (Rabbit anti-Lactoferrin (006) [HRP] (Novus; Cat: NBP2-89794H)) was added and incubated at room temperature for 1 hour. Plates were washed and then developed by adding Slow Step ELISA TMB substrate (Thermofisher, Cat: 34024) to each well. Plates were incubated for 12 minutes, and 2.0M aqueous sulfuric acid was added to quench the reaction. Absorbance was measured using a plate reader; at 450 nanometers and 652 nanometers (Spectramax M2). Lactoferrin concentration was interpolated from the standard curve; limits of quantitation were 36-2100 ng/mL.

#### Bovine Lactoferrin ELISA

A Sandwich ELISA (Bovine Lactoferrin ELISA kit, NBP3-12185, Novus Biologicals) was used and adopted to determine bovine lactoferrin concentrations in rat serum. This assay was modified to detect bovine lactoferrin ranging from 6.9 – 5000 ng/ml. The assay was performed per the manufacturer’s instructions with the following modifications: standards were prepared using bovine LF (Lactoferrin Company; Product 11683) that was used in the oral administration to rats, diluted in 1:10 rat serum, positive controls were prepared by spiking known concentrations of bovine LF into 1:10 rat serum, and the study samples were diluted 1:10 prior to testing. Lactoferrin concentration was interpolated from the standard curve; limit of quantitation was 69-50,000 ng/mL.

### Immunophenotyping

After terminal necropsy, spleen samples were collected from the main study animals and three toxicokinetic animals/sex/group on Day 15, blind coded, stored refrigerated (2-8℃) and shipped to the test site at Michigan State University (East Lansing, MI) for exploratory spleen immunophenotyping analysis to determine whether oral administration of rhLF protein to rats produce changes in the frequency of major immune cell types in the spleen as evaluated by flow cytometry. Immunophenotyping was conducted blinded to dose level.

#### Tissue processing

Splenocytes were isolated by mechanical disruption of spleen samples and made into single cell suspensions in RPMI 1640 medium supplemented with 5% FCS and penicillin/streptomycin. Red blood cells were lysed using Zap-oglobin-II Lytic Reagent (Beckman Coulter) prior to counting splenocytes on a Z1 Beckman Coulter Counter.

#### Flow Cytometry

A total of 2 x 10^6^ splenocytes were placed in a 96-well round bottom culture plate for surface and intracellular antibody staining. Splenocytes were washed using Hank’s Balanced Salt Solution (HBSS, (pH 7.4)), (Invitrogen) and stained with LIVE/DEAD Fixable Near-IR Dead Cell Stain (Gibco Invitrogen) to assess cell viability. Splenocytes were washed with FACS buffer (1× HBSS (pH 7.5) containing 1% BSA and 0.1% sodium azide) and Cell FcRs were blocked with purified mouse anti-rat CD32 (BD Biosciences). After a 15-minute incubation at 4°C, splenocytes were stained for surface proteins.

For the Granulocyte/Macrophage/DC and B cell panel: CD45 (OX-1) (Biolegend), CD172a (clone OX-41) (BD Bioscience), CD11b (clone WT.5) (Biolegend), CD25 (clone OX-39) (Biolegend), and AffiniPure Anti-Rat IgG + IgM (Jackson ImmunoResearch). For the T cell panel: CD45 (OX-1) (Biolegend), CD3 (clone 1F4) (Biolegend), CD4 (clone W3/25) (Biolegend), CD8 (clone OX-8) (Biolegend) and CD25 (clone OX-39) (Biolegend).

For FoxP3+ T cells: Splenocytes were incubated at 4°C and washed three times with FACS buffer and fixed with FoxP3/Transcription Factor Staining Buffer Set (Invitrogen) fixative and stored at 4°C until intracellular staining. To measure intracellular FoxP3, splenocytes were washed and incubated with 1x Permeabilization buffer solution (Invitrogen) for 15 minutes, then incubated with anti-rat FoxP3 (Invitrogen) for 30 minutes. Splenocytes were washed 3X with the permeabilization buffer and then washed one more time with FACS buffer and resuspended in FACS buffer.

Flow cytometric analysis was performed on a Cytek Northern Lights full spectral analyzer (Cytek Biosciences) and analyzed using FlowJo v10.9.0 (Tree Star, Ashland, OR) software.

### Statistical analysis

#### In-life, clinical pathology, and postmortem measures

Statistical analyses for in-life, clinical pathology, and postmortem measures were conducted with Provantis version 9. Raw data was tabulated within each time interval, and the mean and standard deviation and/or incidence counts (categorical variables) were calculated for each endpoint by sex and group. For each endpoint, treatment groups were compared to the control group. The groups were compared using an overall one-way ANOVA F-test if Levene’s test was not significant, or the Kruskal-Wallis test if it was significant. If the overall F-test or Kruskal-Wallis test was found to be significant, then pairwise comparisons were conducted using Dunnett’s or Dunn’s test, respectively. Results of all pairwise comparisons were reported at the 0.05 and 0.01 significance levels and all endpoints were analyzed using two-tailed tests unless indicated otherwise.

#### Immunophenotyping

Statistical analyses were performed using GraphPad Prism version 10.0.02 (GraphPad Software, La Jolla, CA). To determine statistically significant changes between the treatment groups and the control (0 mg/kg/day) within male and female rats, a one-way ANOVA with a Dunnet’s multiple comparisons posttest was used. The mean ± SD is displayed in all bar graphs (* p = < 0.05, ** p ≤ 0.01, *** p ≤ 0.001, and **** p ≤ 0.0001).

## RESULTS

### Ingestion of Helaina rhLF was well tolerated up to 2000 mg/kg/day

All animals survived the 14-day oral dosing. There was no morbidity or mortality associated with administration of rhLF or bLF and in-life observations showed minimal instances of abnormal physical signs, none that were attributed to treatment with rhLF or bLF. Changes in body weight during the study are shown in Figure 1. No significant body weight changes were observed during the course of treatment for any dose level. At necropsy, there were no changes attributable to the administration of test articles in males or females, and there were no significant differences in absolute organ weights or the relative organ weights between male or female control and dosed groups. Any observed differences were not statistically significant, were not clearly dose-responsive in nature, and/or were generally due primarily to differences for an individual animal. No macroscopic observations were noted at any dose level evaluated, including controls. Collectively, these data show that ingestion of rhLF and bLF is well tolerated up to at least 2000 mg/kg/day.

**Figure 1:**
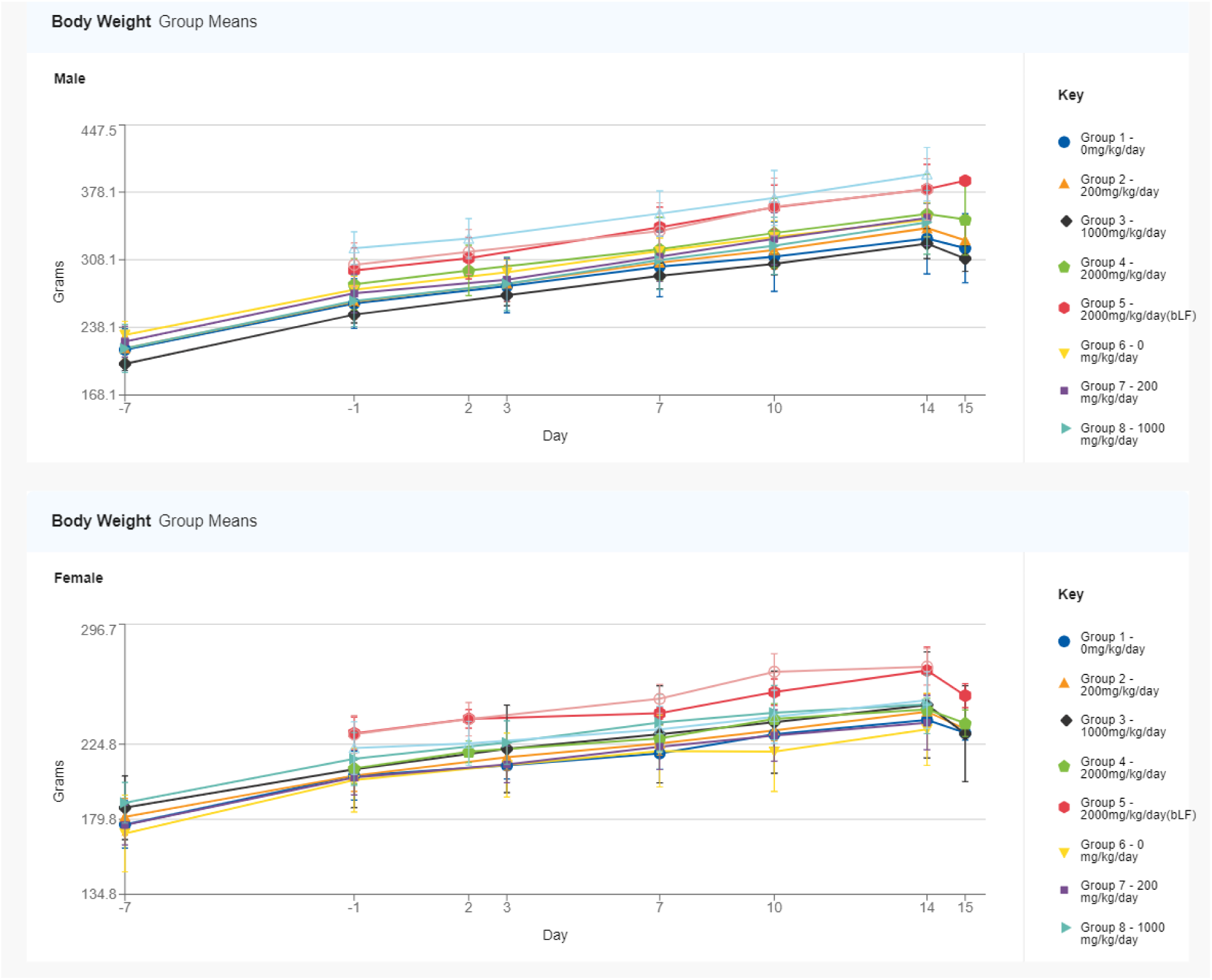
Body Weight Changes by Sex. Body weight was measured bi-weekly for each animal during the treatment period for both main (Groups 1-5) and LF absorption animals (Group 6-10) and the mean results shown over time. Males are shown on the top figure and females on the bottom figure.

### Minimal to moderate changes in hematology and chemistry parameters were observed at ≥1000 mg/kg/day

Recombinant hLF and bLF-related alterations in measured hematology parameters were observed in both sexes at 2000 mg/kg/day and are displayed in Table 1. These alterations comprised higher hemoglobin distribution width (HDW), higher mean platelet volume (MPV), lower platelet distribution width (PDW), lower mean platelet component (MPC), and/or lower platelet component distribution width (PCDW). At termination on Day 15, in females at 2000 mg/kg/day rhLF and 2000 mg/kg/day bLF, there were test article-related, minimally higher HDW, reflecting greater variation in erythrocyte hemoglobin content. In both sexes at 2000 mg/kg/day rhLF and 2000 mg/kg/day bLF, there were test article-related, minimally to mildly higher MPV, reflecting larger mean platelet size. Additionally, females at 2000 mg/kg/day rhLF and 2000 mg/kg/day bLF exhibited test article-related, minimally lower PDW, reflecting less variation in platelet size. In females at 2000 mg/kg/day rhLF and 2000 mg/kg/day bLF, there were test article-related, minimally lower MPC values, reflecting lower platelet refractive index, or lower platelet density / granularity; additionally, both sexes at 2000 mg/kg/day rhLF and 2000 mg/kg/day bLF exhibited test article-related, minimally to mildly lower PCDW, reflecting less variation in platelet density / granularity.

**Table 1:**
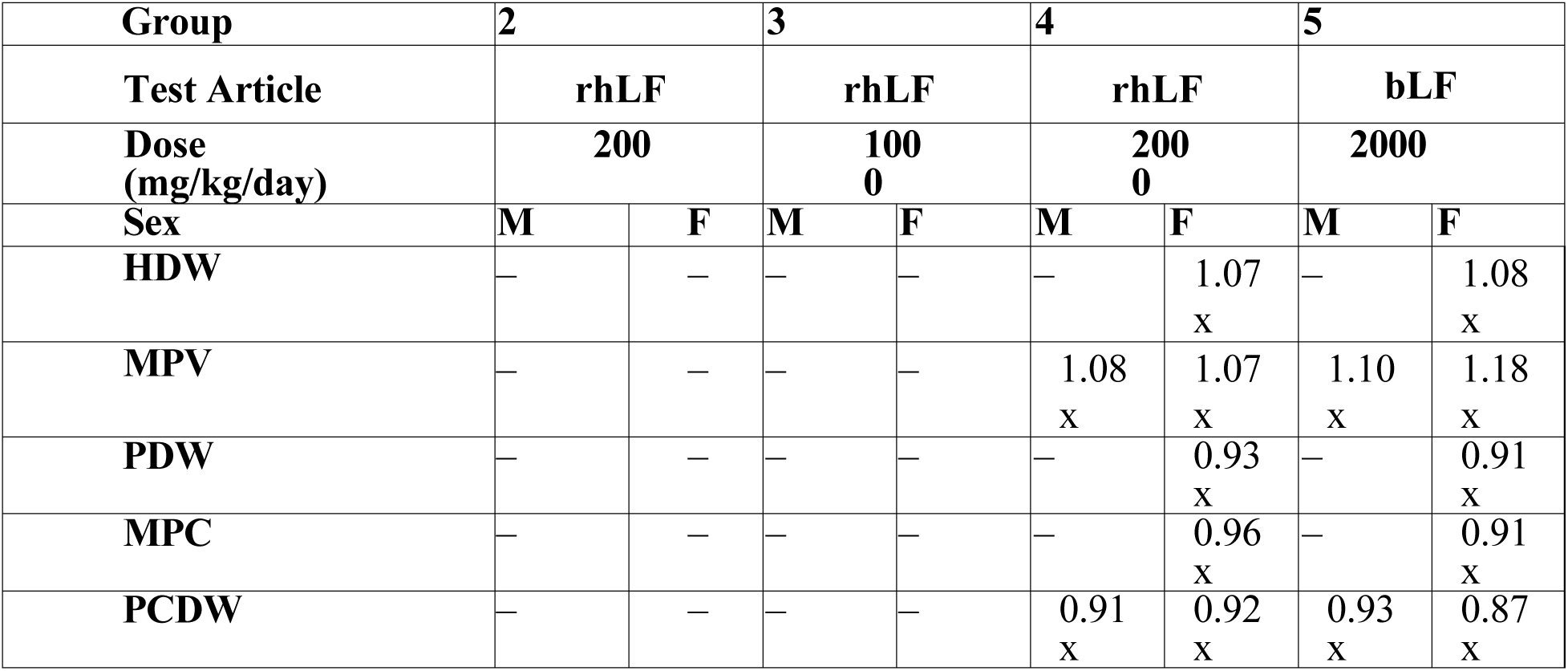
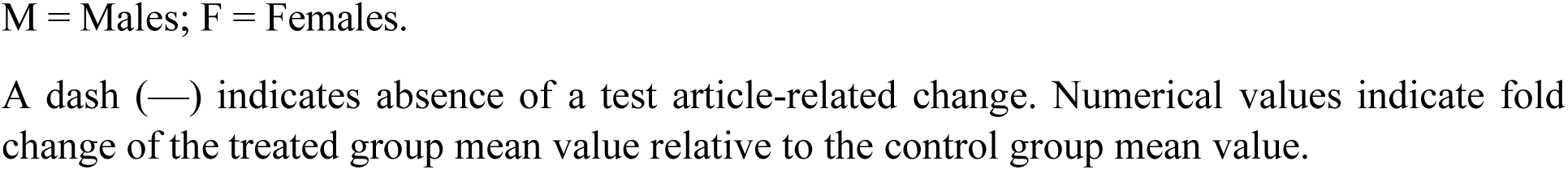
rhLF– and bLF-Related Hematology Changes – Day 15.

| Group | 2 |  | 3 |  | 4 |  | 5 |  |
| --- | --- | --- | --- | --- | --- | --- | --- | --- |
| Test Article | rhLF |  | rhLF |  | rhLF |  | bLF |  |
| Dose (mg/kg/day) | 200 |  | 100<br>0 |  | 200<br>0 |  | 2000 |  |
| Sex | M | F | M | F | M | F | M | F |
| HDW | — | — | — | — | — | 1.07<br>x | — | 1.08<br>x |
| MPV | — | — | — | — | 1.08<br>x | 1.07<br>x | 1.10<br>x | 1.18<br>x |
| PDW | — | — | — | — | — | 0.93<br>x | — | 0.91<br>x |
| MPC | — | — | — | — | — | 0.96<br>x | — | 0.91<br>x |
| PCDW | — | — | — | — | 0.91<br>x | 0.92<br>x | 0.93<br>x | 0.87<br>x |

M = Males; F = Females.
A dash (—) indicates absence of a test article-related change. Numerical values indicate fold change of the treated group mean value relative to the control group mean value.

Helaina rhLF-related alterations in measured clinical chemistry parameters were observed in males at 2000 mg/kg/day and females at ≥ 1000 mg/kg/day, and bLF-related alterations in measured clinical chemistry parameters were observed in both sexes at 2000 mg/kg/day, as shown in Table 2. At termination on Day 15, in both sexes at 2000 mg/kg/day rhLF and males at 2000 mg/kg/day bLF, there were test article-related, minimally higher total bilirubin (TBIL) concentrations. In males at 2000 mg/kg/day rhLF and in females at 2000 mg/kg/day bLF, there were test article-related, minimally higher urea nitrogen (UREAN) concentrations. In both sexes at 2000 mg/kg/day rhLF, there were test article-related, minimally lower mean creatinine (CREAT) concentrations. In males at 2000 mg/kg/day rhLF, there were test article-related, minimally to mildly higher cholesterol (CHOL) concentrations. Two individual females at 2000 mg/kg/day rhLF and one male and two females at 2000 mg/kg/day bLF also exhibited minimally to mildly higher CHOL concentrations which were potentially test article-related, though the small magnitude of difference and inconsistent effect among cohort group members precluded definitive interpretation. In females at 2000 mg/kg/day rhLF and 2000 mg/kg/day bLF, there were test article-related, mildly higher triglyceride (TRIG) concentrations. One male at 2000 mg/kg/day rhLF and one male at 2000 mg/kg/day bLF also exhibited mildly higher TRIG concentrations which were potentially test article-related, though the small magnitude of difference, inconsistent effect among cohort group members, and high variability inherent to TRIG concentration precluded definitive interpretation. In both sexes at 2000 mg/kg/day rhLF and 2000 mg/kg/day bLF, there were test article-related, minimally to mildly higher albumin (ALB) concentrations with resultant higher A:G ratios. In males at 2000 mg/kg/day and in females at ≥ 1000 mg/kg/day rhLF, and in both sexes at 2000 mg/kg/day bLF, there were test article-related, minimally to mildly lower mean phosphorus (PHOS) concentrations. In both sexes at 2000 mg/kg/day rhLF and females at 2000 mg/kg/day bLF, there were test article-related, minimally lower sodium (NA) and/or chloride (CL) concentrations.

**Table 2:**
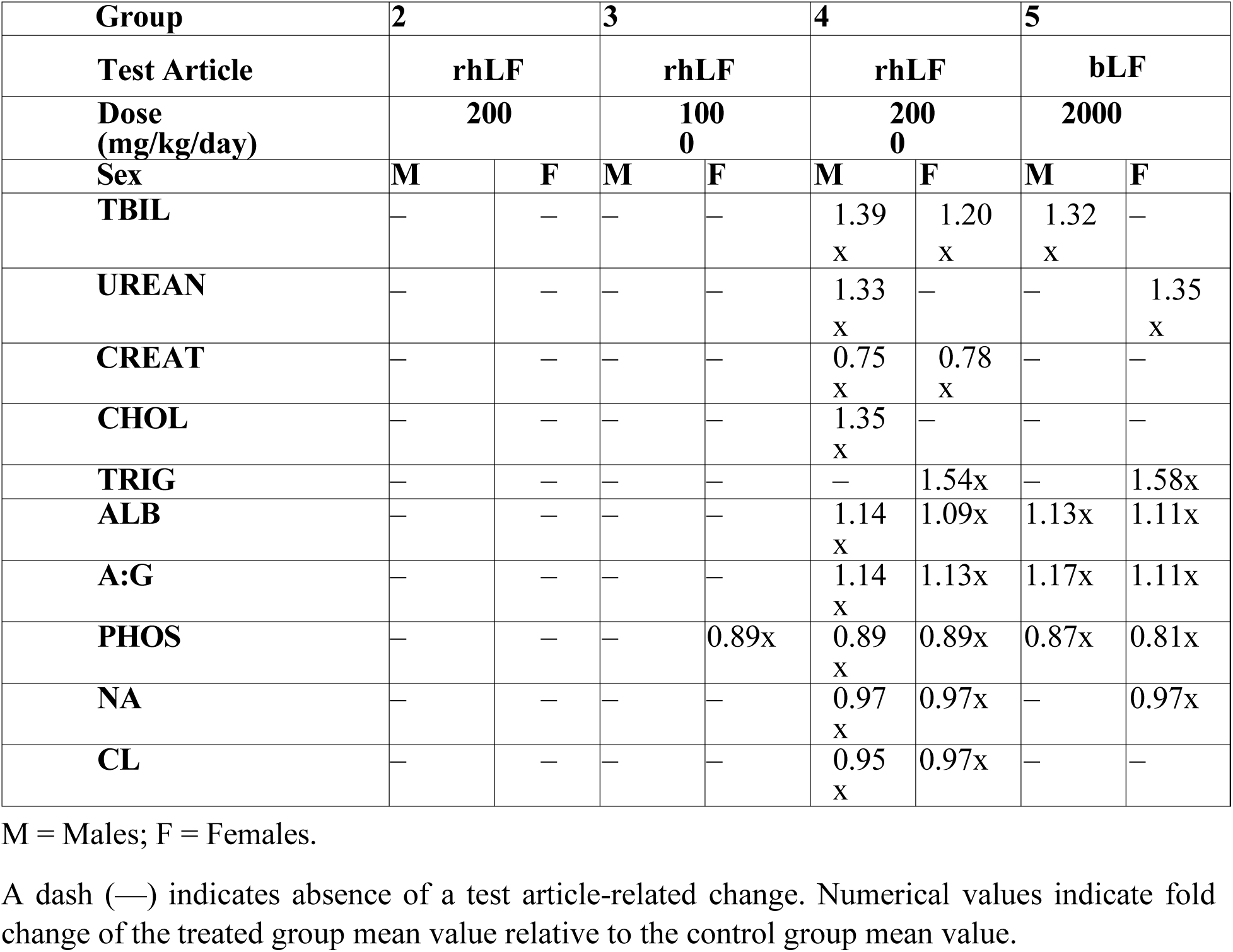
rhLF– and bLF-Related Clinical Chemistry Changes – Day 15.

| Group | 2 |  | 3 |  | 4 |  | 5 |  |
| --- | --- | --- | --- | --- | --- | --- | --- | --- |
| Test Article | rhLF |  | rhLF |  | rhLF |  | bLF |  |
| Dose (mg/kg/day) | 200 |  | 100<br>0 |  | 200<br>0 |  | 2000 |  |
| Sex | M | F | M | F | M | F | M | F |
| TBIL | — | — | — | — | 1.39<br>x | 1.20<br>x | 1.32<br>x | — |
| UREAN | — | — | — | — | 1.33<br>x | — | — | 1.35<br>x |
| CREAT | — | — | — | — | 0.75<br>x | 0.78<br>x | — | — |
| CHOL | — | — | — | — | 1.35<br>x | — | — | — |
| TRIG | — | — | — | — | — | 1.54x | — | 1.58x |
| ALB | — | — | — | — | 1.14<br>x | 1.09x | 1.13x | 1.11x |
| A:G | — | — | — | — | 1.14<br>x | 1.13x | 1.17x | 1.11x |
| PHOS | — | — | — | 0.89x | 0.89<br>x | 0.89x | 0.87x | 0.81x |
| NA | — | — | — | — | 0.97<br>x | 0.97x | — | 0.97x |
| CL | — | — | — | — | 0.95<br>x | 0.97x | — | — |

M = Males; F = Females.
A dash (—) indicates absence of a test article-related change. Numerical values indicate fold change of the treated group mean value relative to the control group mean value.

In terms of iron-related parameters (Table 3), on Day 15, in males at 2000 mg/kg/day rhLF and 2000 mg/kg/day bLF, there were mildly to moderately lower ferritin (FERR) concentrations, suggesting lower quantities of stored iron; conversely, there were concurrent minimally to mildly higher serum iron (FE) and total iron binding capacity (TIBC) concentrations and transferrin saturation (TFRRNSAT) values. These parameters tended to be highly variable among females and lacked a clear test article-related pattern of results.

**Table 3:**
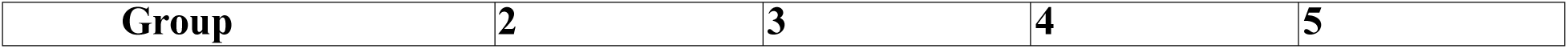
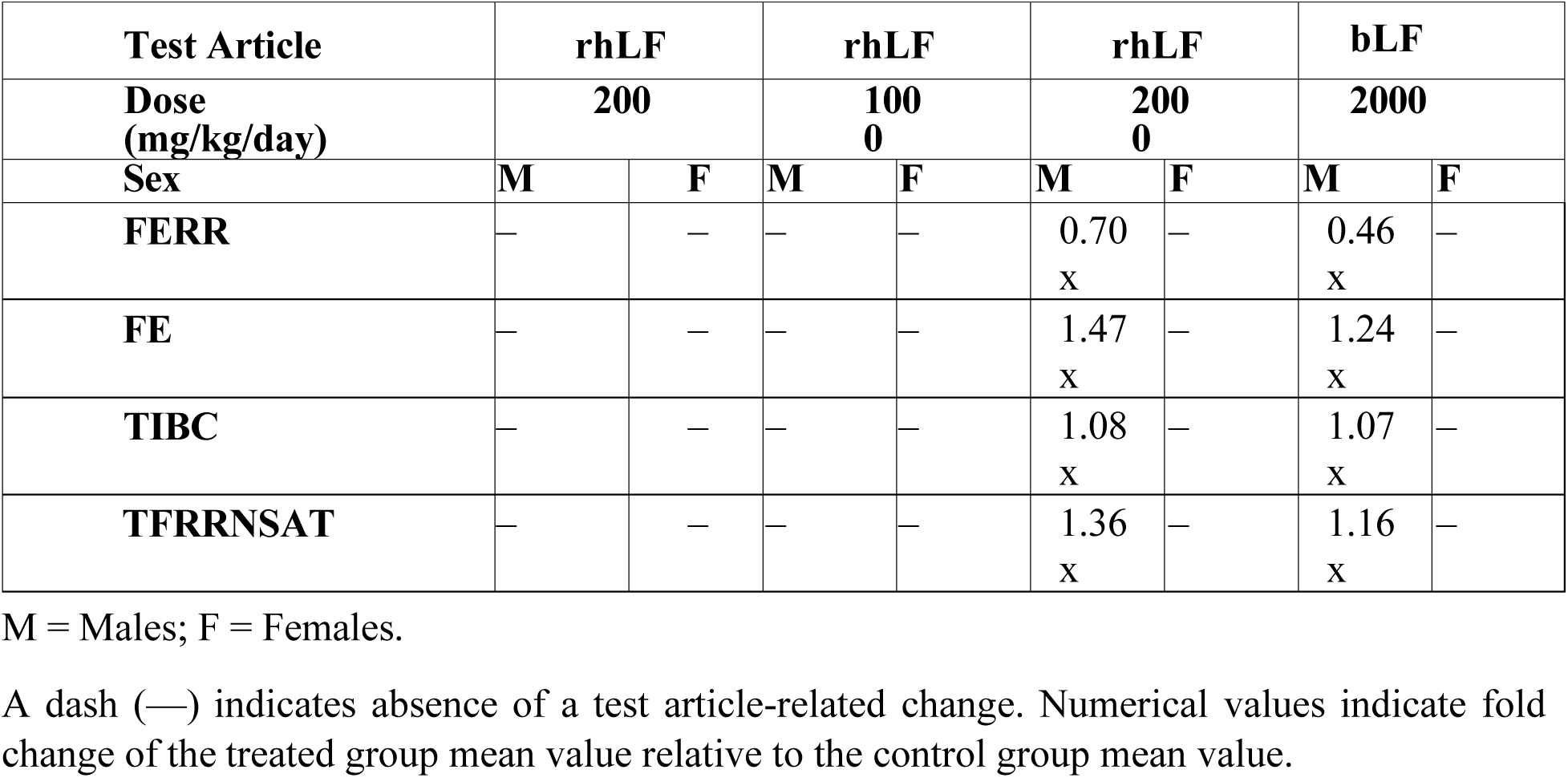
rhLF– and bLF-Related Clinical Chemistry Changes – Iron-Related Parameters-Day 15.

| **Test Article** | **rhLF** |  | **rhLF** |  | **rhLF** |  | **bLF** |  |
| --- | --- | --- | --- | --- | --- | --- | --- | --- |
| **Dose**<br>**(mg/kg/day)** | **200** |  | **100**<br>**0** |  | **200**<br>**0** |  | **2000** |  |
| **Sex** | **M** | **F** | **M** | **F** | **M** | **F** | **M** | **F** |
| **FERR** | — | — | — | — | 0.70<br>x | — | 0.46<br>x | — |
| **FE** | — | — | — | — | 1.47<br>x | — | 1.24<br>x | — |
| **TIBC** | — | — | — | — | 1.08<br>x | — | 1.07<br>x | — |
| **TFRNSAT** | — | — | — | — | 1.36<br>x | — | 1.16<br>x | — |

M = Males; F = Females.
A dash (—) indicates absence of a test article-related change. Numerical values indicate fold change of the treated group mean value relative to the control group mean value.

Remaining fluctuations among individual and mean values, regardless of statistical significance, were considered sporadic, consistent with biologic variation, and/or negligible in magnitude, and not related to test article administration. Overall, the clinical pathology changes that were observed in rhLF were also observed in bLF, suggesting a similar toxicological profile between the two proteins.

### Ingestion of Helaina rhLF or bLF leads to minimal alterations in immune cell phenotypes within the spleen

Ex vivo immunophenotyping techniques using immune organs have been used as a measure of immunotoxicity related to test article administration.^30^ To determine whether ingestion of rhLF has an impact on immune cell types within the spleen, immunophenotyping via flow cytometry was performed after 14 days of continuous exposure and the gating strategy is shown in Supplemental Figure 1A. The spleens were collected following euthanasia and dispersed into single cell suspension for flow cytometry staining and first analyzed for cell viability. It was determined that oral gavage with Helaina rhLF or with bLF did not have any significant effects on cell viability in either female or male rats in any of the treatment groups (Supplemental Figure 1B).

Next, spleen resident immune cells were assessed for the myeloid marker, CD172a^+^, expression, and there was a significant, and similar, increase in the frequency of immune cells expressing CD172a^+^ in both male and female rats administered the highest dose (2,000 mg/kg/day) of either rhLF or bLF (Figure 2A). While there was a significant increase in the percentage of CD172a^+^ spleen cells in response to high dose LF (i.e., rhLF and bLF), there was no change in the percentage of cells expressing CD11b (Supplemental Figure 1C), suggesting that the change in CD172a^+^ cells was not due to an increase in monocyte or macrophage cells. While treatment with high doses of LF did not affect the frequency of monocytes/macrophages, when the percentage of CD172a^+^ cells expressing CD11b and CD25, a marker of dendritic cells, was assessed, a significant increase was found at the 2,000 mg/kg/day dose group (Figure 2B). This would suggest that most of the changes in the CD172a^+^ expressing cell population are due to the observed increase in DC. The final immune cell type evaluated, via use of antibody panel 1, (Supplemental Figure 1A) were B cells as identified by the presence of B cell receptors on the cell surface plasma membrane (IgM^+^ or IgG^+^). When the frequency of B cells in the spleen were quantified, no treatment-related changes were observed in either female or male rats following oral gavage with LF protein (Figure 2C).

**Figure 2.**
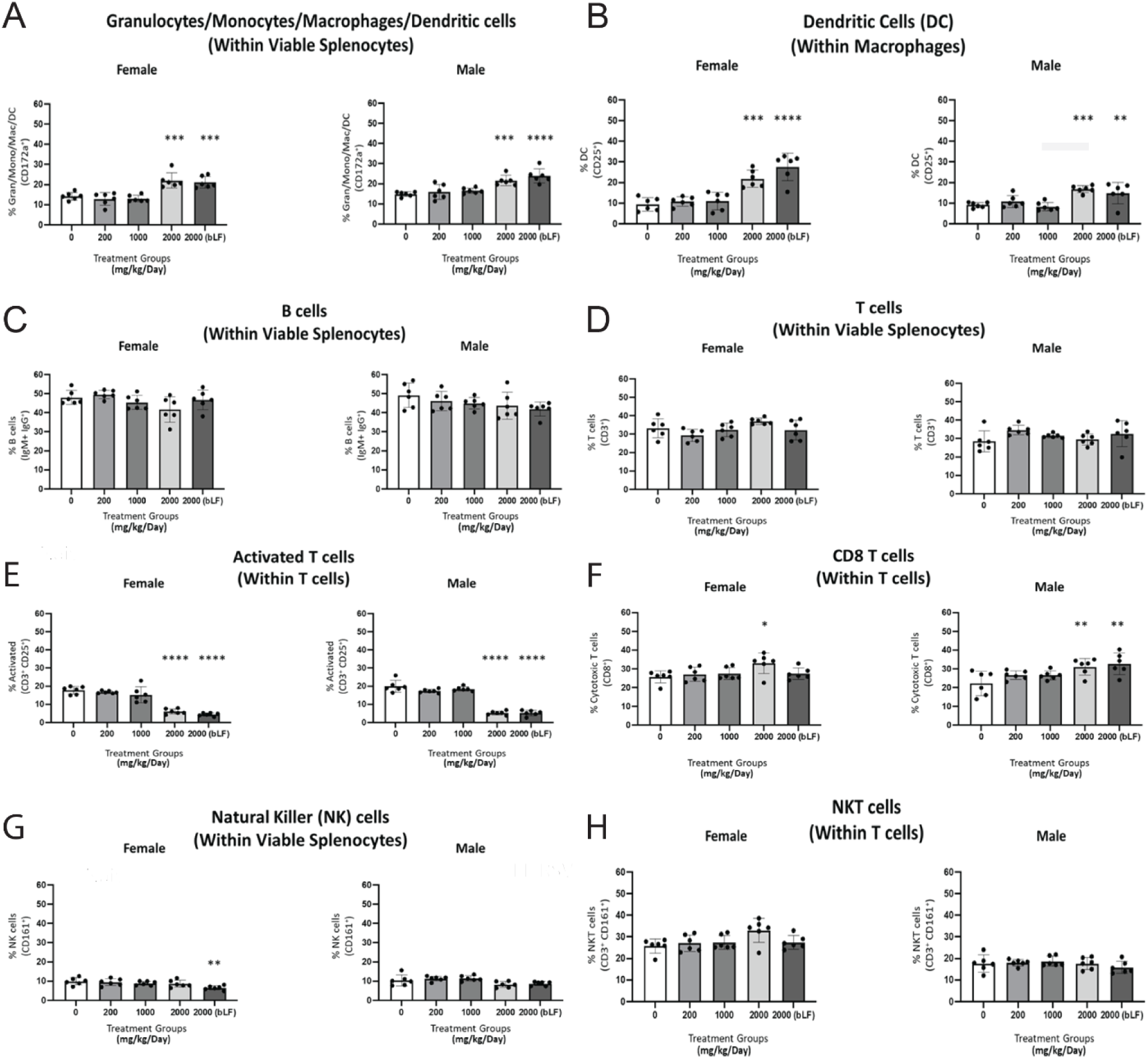
Ingestion of Helaina rhLF or bLF Leads to Minimal Alterations in Immune Cell Phenotypes within the Spleen. Spleens were collected after the 14-day exposure period and dispersed into single-cell suspension for immunophenotyping analysis via flow cytometry. All cells were gated within the live cell gate within the splenocyte population. **A**. CD172a+ myeloid-type cells (granulocytes, monocytes, macrophages, dendritic cells). **B**. CD25+ Dendritic cells (within CD17a+/CD11b+ population). **C**. IgM+/IgG+ B cells (within the splenocyte population). **D**. CD3+ T cells (within splenocyte population). **E**. CD25+ Activated T cells (within CD3+ T cell population). **F**. CD8+ T cells (within CD3+ T cell population). **G**. CD161+ natural killer (NK) cells (within splenocyte population). **H**. CD3+CD161+ NKT cells (within CD3+ T cell population). N = 6 animals per group. * = p < 0.05, ** = p ≤ 0.01, *** = p ≤ 0.001, **** = p ≤ 0.0001

Antibody panel 2 (Supplemental Figure 1A) was used to quantify the frequency of T lymphocyte populations in the spleen of rats gavaged with either rhLF or bLF. There were no treatment-mediated changes in the frequency of immune cells in the spleen expressing CD3, which is indicative of total splenic T cells (Figure 2D). However, when the percentage of activated T cells was assessed by measuring the percentage of CD25^+^ CD3^+^ T cells, there was a marked decrease in the frequency of these cells within the spleen at the highest dose (2,000 mg/kg/day) of either rhLF or bLF (Figure 2E). Despite observing no effect on the total frequency of CD3^+^ cells in the spleen in any of the treatment groups, when the populations of T cells were evaluated, there were treatment-related effects at the population level. For example, when the frequencies of CD8^+^ T cells within the spleen were assessed, we found that the highest dose treatment with rhLF resulted in statistically significant increases in the percentage of CD8^+^ T cells in both female and male rats, and an increase in male rats given bLF (Figure 2F). This increase was not observed in the frequencies of spleen resident CD4^+^ T cells as there was no change in any of the treatment groups in either male or female rats (Supplemental Figure 1D). We next assessed a minor population of T cells with regulatory function (Treg), which are typically induced in response to inflammation and T cell activation. These cells are identified within the CD4^+^ T cell compartment by their expression of CD25 and the intracellular protein FoxP3. No significant changes in the frequency of Treg cells was observed in any of the treatment groups (Supplemental Figure 1E).

Finally, we examined the effects of oral administration of LF protein on the frequencies of natural killer (NK) cells as evidenced by changes in the expression of CD161^+^ (Supplemental Figure 1A). Recombinant hLF administration did not produce any changes in the frequency of CD161^+^ cells. By contrast, administration of bLF (2000 mg/kg/day) produced a modest reduction in the frequency of CD161^+^ cells in female, but not male rats (Figure 2G). We then examined a subpopulation of NK cells, NK T cells, which are marked by the expression of CD3. There were no treatment-related changes in either male or female rats in the frequencies of spleen resident NK T cells (Figure 2H).

### Postprandial levels of LF detected in the serum

In order to assess potential bioavailability of rhLF and bLF in circulation, serum was collected at multiple time points (0.5, 1, 3, 6, 12, 24 hours) following LF administration on Days 1 and 14 and tested for the presence of LF using ELISA. Results indicate that rhLF is quickly absorbed and detected in serum within 30 minutes of administration for males dosed at 2000 mg/kg/day on both Day 1 and Day 14 (Figure 3A, B), and very quickly cleared from circulation with no detection observed by 3 hours post-administration (Figure 3A,B). A similar trend was observed in female rats administered 2000 mg/kg/day rhLF on Day 1 (Figure 3C). However, on Day 14, female rats administered ≥ 1000 mg/kg/day had detectable levels of rhLF that cleared circulation by 3-6 hours post-administration (Figure 3D). No bLF was detected in the serum from male rats administered 2000 mg/kg/day (Figure 3A,B), and very low levels of bLF was detected in female rats (Figure 3C,D). These data suggest that Helaina rhLF may be more bioavailable than bLF with a stronger response in females; however, a lack of detectable LF in most samples tested and high variability amongst responders make it difficult to fully assess.

**Figure 3.**
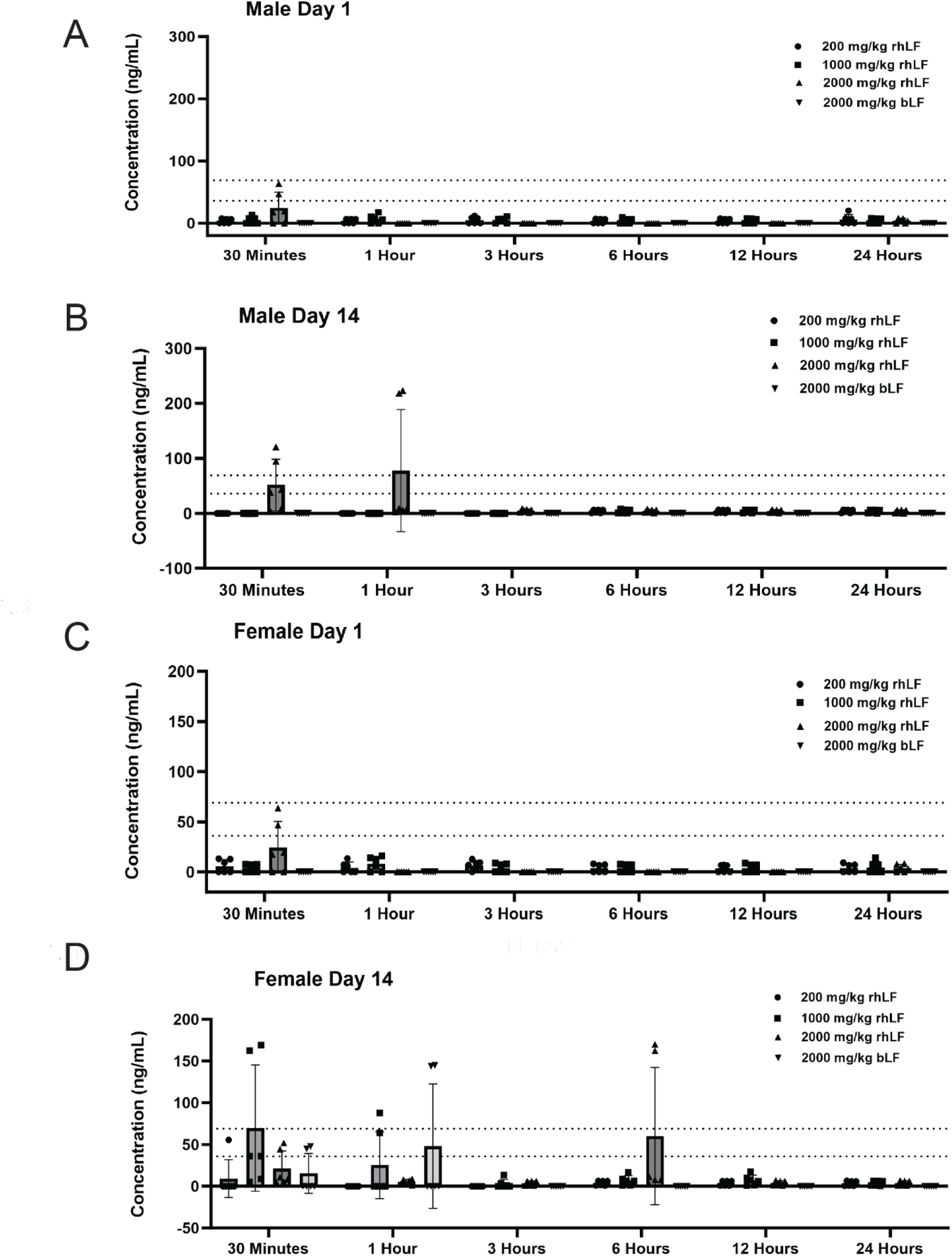
Postprandial levels of LF detected in the serum. Serum was collected over multiple time points after administration on Day 1 and Day 14 of dosing. ELISA was performed to detect LF (rhLF or bLF) in the serum. **A**. Day 1 LF results in male rats **B**. Day 14 LF results in male rats. **C**. Day 1 LF results in female rats. **D**. Day 14 results in female rats. N = 3 animals per time point. No statistical analysis was performed due to insufficient data/lack of responses above the limit of detection of the assays.

## DISCUSSION

To date, no published study has evaluated the safety and oral toxicity of the novel protein, Helaina rhLF, produced in *K. phaffii.* Currently, the only LF commonly used for commercial food applications (e.g., added to yogurt and consumed as a dietary supplement) is bLF. The present study aimed to assess the oral toxicity of Helaina rhLF compared to bLF administration in adult Sprague Dawley rats by once daily oral gavage for 14 days and was designed as a dose range-finding study to inform dosages for future definitive studies. Results showed that Helaina rhLF was well tolerated in rats up to 2000 mg/kg/day, or 400x Helaina’s intended commercial use (0.34 g/day in an average 70 kg adult). All animals survived to terminal necropsy. There were no test article-related effects on clinical observations, body weights, organ weights or macroscopic observations.

Changes in hematology parameters (increased HDW, higher MPV, lower PDW, lower MPC, and/or lower PCDW) were limited to the 2000 mg/kg groups and were similar in both sexes. Minimal to moderate rhLF-related alterations in measured clinical chemistry parameters (higher TBIL, UREAN, CHOL or TRIG, ALB, A:G ratios, and lower CREAT, PHOS, NA and/or CL concentrations) were observed in males at 2000 mg/kg/day and females at ≥ 1000 mg/kg/day, and bLF-related alterations in measured clinical chemistry parameters were observed in both sexes at 2000 mg/kg/day. While these changes were considered statistically significant and biologically relevant, the consequences of these changes are unknown without corroborative histopathology data. However, it is important to note that these changes did not outwardly affect the health and well-being of the animals and support the use of 2000 mg/kg/day as a high dose for modeling toxicity by producing some toxic effects, but not excessive lethality that would prevent meaningful evaluation.

Lactoferrin is an iron-binding protein^36^; therefore, this study also evaluated parameters essential for iron homeostasis. Changes in these parameters were observed in males at 2000 mg/kg/day and females at ≥ 1000 mg/kg/day, as well as bLF-treated groups in both sexes at 2000 mg/kg/day. Results included lower ferritin concentrations, higher serum iron concentrations, higher total iron binding capacity, and/or higher transferrin saturation. These changes could indicate that iron is being transported out of the cells and into circulation, and correlates with clinical studies using bLF to treat inflammatory-driven anemia in pregnant and non-pregnant women.^12,13^ It is important to note that while Helaina rhLF is greater than 50% saturated with iron, the bovine LF used in these studies was less than 10% iron saturated, indicating that inherent iron saturation status is not a pivotal indicator for iron homeostasis properties of LF.

The present work also determined whether oral administration of rhLF produces changes in the frequency of major immune cell types in the spleen as evaluated by flow cytometry as a measure of potential immunotoxicity.^28–30^ Collectively, this study shows an absence of any changes in the frequency of major immunocompetent cell populations in the spleen, comprising both myeloid and lymphoid lineages, after administration of rhLF at doses below 2000 mg/kg/day. Modest changes in the frequency of several spleen cell populations, including increases in CD172a^+^ DC, and CD8^+^ T cells, and decreases in activated CD25^+^ T cells, were observed in both rhLF and bLF treatment groups, which was restricted to the highest dose group (2000 mg/kg/day). Predominantly, these changes in cell population frequency occurred in both male and female rats, suggesting no sex-specific alterations. Further, these results suggest that there are no differences between rhLF and bLF in inducing immunological changes in the Sprague Dawley rat. These changes observed in spleen cell type frequencies in both rhLF and bLF at 2,000 mg/kg/day treatment groups can neither be deemed detrimental nor beneficial, as no other measurements of the immune system were conducted in this analysis and should be considered in future safety studies.

Assessment of LF absorption was performed at multiple time points ranging from 30 minutes to 24 hours and indicated very low levels of LF absorption overall. Males were observed to have lower levels of both rhLF and bLF than females, indicating a potential sex difference; however, future studies should include additional time points to better assess these findings. Importantly, none of the male rats had detectable levels of bLF in the serum, and detectable levels of bLF were only observed in three female samples, suggesting better bioavailability of rhLF compared to bLF. Additionally, it was observed that LF was quickly absorbed into the serum (average 30 minutes to one hour) and quickly cleared from circulation (no detectable levels after ∼3 hours), with no signs of accumulation. These findings are correlated with the literature that identified LF in circulation within 10-20 minutes after ingestion in a mouse model^37^ and within ∼30 minutes in humans.^38^ Finally, studies that injected LF directly into the blood via intravenous injection showed the half-life of LF was 10 minutes in adult rats^39^ respectively, with complete removal by 7 hours after injection,^40^ supporting these findings of clearance.

In conclusion, administration of Effera™ (Helaina rhLF) by once daily oral gavage for 14 days was well tolerated in rats and corroborates previous general toxicity studies showing administration of high levels of rhLF (≤ 2000 mg/kg/day) are safe and tolerable in the in vivo rat system.^23,24^ These studies also showed similar toxicological effects between rhLF and bLF, supporting the safe use of rhLF into foods that currently contain bLF. Additionally, this study has informed the future definitive toxicology studies, indicating that these studies should include doses of rhLF up to 2000 mg/kg/day to allow for proper toxicological assessment.

## Abbreviations

*K. phaffii*: Komagataella phaffii
LF: Lactoferrin
rhLF: Recombinant human lactoferrin
bLF: Bovine lactoferrin
hLF: Human lactoferrin
hmLF: Human milk lactoferrin
GRAS: Generally Recognized as Safe
NOAEL: No observed adverse effect level
HPLC: High performance liquid chromatography
SDS-PAGE: Sodium dodecyl sulfate–polyacrylamide gel electrophoresis
MPV: Mean platelet volume
CHCM: Cellular hemoglobin concentration mean
PDW: Platelet distribution width
PCT: Platelitcrit
CH: Mean cellular hemoglobin content
MCH: Mean corpuscular volume
MCHC: Mean corpuscular hemoglobin concentration (calculated)
RDW: Red blood cell distribution width
HDW: Hemoglobin distribution width
PCDW: Platelet component distribution width
CHr: Reticulocyte hemoglobin content
MCVr: Mean reticulocyte volume
UIBC: Unsaturated iron binding capacity
TIBC: Total iron binding capacity
TSAT: Transferrin saturation
TBIL: Total bilirubin
UREAN: Urea nitrogen
CREAT: Creatinine
CHOL: Cholesterol
TRIG: Triglycerides
ALB: Albumin
PHOS: Phosphorus
NA: Sodium
CL: Chloride
FERR: Ferritin
FE: Iron
TIBC: Total iron binding capacity
TFRRNSAT: Transferrin saturation

## ACKNOWLEDGEMENTS

The authors would like to thank technical writer Sarah M. Weyers, PhD, RD, CAPM for assistance with the preparation of the manuscript, Yanisa Anaya, BE and Raysa Rosario Martinez, BS for their technical expertise in lactoferrin ELISA methodology.

## FOOTNOTES

Funding: Privately funded by Helaina, Inc.

Declaration of Conflicting Interest: Authors RP, JSS, BF, RV-D, AJC, and C-AM are all employees of Helaina, Inc.

**Supplemental Figure 1**. Spleens were collected after the 14-day exposure period and dispersed into single-cell suspension for immunophenotyping analysis via flow cytometry. **A**. Overall gating strategy. **B**. Viable cells (within CD45+ splenocyte population). **C**. CD11b+ cells (within CD172a+ population). **D**. CD4+ helper T cells (within CD3+ T cell population). **E**. CD25+FoxP3+ regulatory T cells (within CD4+ T cells). N = 6 animals per group.

## Data Availability

Not applicable

---

**Supplemental Table 1:** Hematologic and Clinical Chemistry Parameters.

| Hematology | Clinical Chemistry (Without Iron) | Clinical Chemistry (Iron Parameters) |
| --- | --- | --- |
| Leukocyte count (total and differential)<br>Erythrocyte count<br>Hemoglobin<br>Hematocrit<br>Absolute reticulocytes<br>Mean platelet volume (MPV)<br>Cellular hemoglobin concentration mean (CHCM)<br>% reticulocytes<br>Platelet distribution width (PDW)<br>Plateletcrit (PCT)<br>Mean cellular hemoglobin content (CH)<br>Mean corpuscular hemoglobin (MCH)<br>Mean corpuscular volume (MCV)<br>Mean corpuscular hemoglobin concentration (calculated) (MCHC))<br>Red blood cell distribution width (RDW)<br>Platelet count | Alkaline phosphatase<br>Total bilirubin<sup>a</sup><br>Aspartate aminotransferase<br>Alanine aminotransferase<br>Urea nitrogen<br>Creatinine<br>Total protein<br>Albumin<br>Globulin<br>Albumin/globulin (A/G) ratio (calculated)<br>Glucose<br>Total cholesterol<br>Triglycerides<br>Electrolytes (sodium, potassium, chloride)<br>Calcium<br>Phosphorus<br>Sample quality | Unsaturated iron binding capacity (UIBC)<br>Total iron binding capacity (TIBC) calculated<br>Iron<br>Ferritin<br>Transferrin saturation (TSAT) |
| • Blood smear (preserve and stain)<br>• Hemoglobin distribution width (HDW)<br>• Platelet component distribution width (PCDW)<br>• Reticulocyte hemoglobin content (CHr)<br>• Mean reticulocyte volume (MCVr) |  |  |

\*When total bilirubin was >0.5 mg/dL, direct bilirubin was also measured, and indirect bilirubin was calculated.

