## Supplemental Figure 1 for "Dose range-finding toxicity study in rats with recombinant human lactoferrin produced in *Komagataella phaffii*"

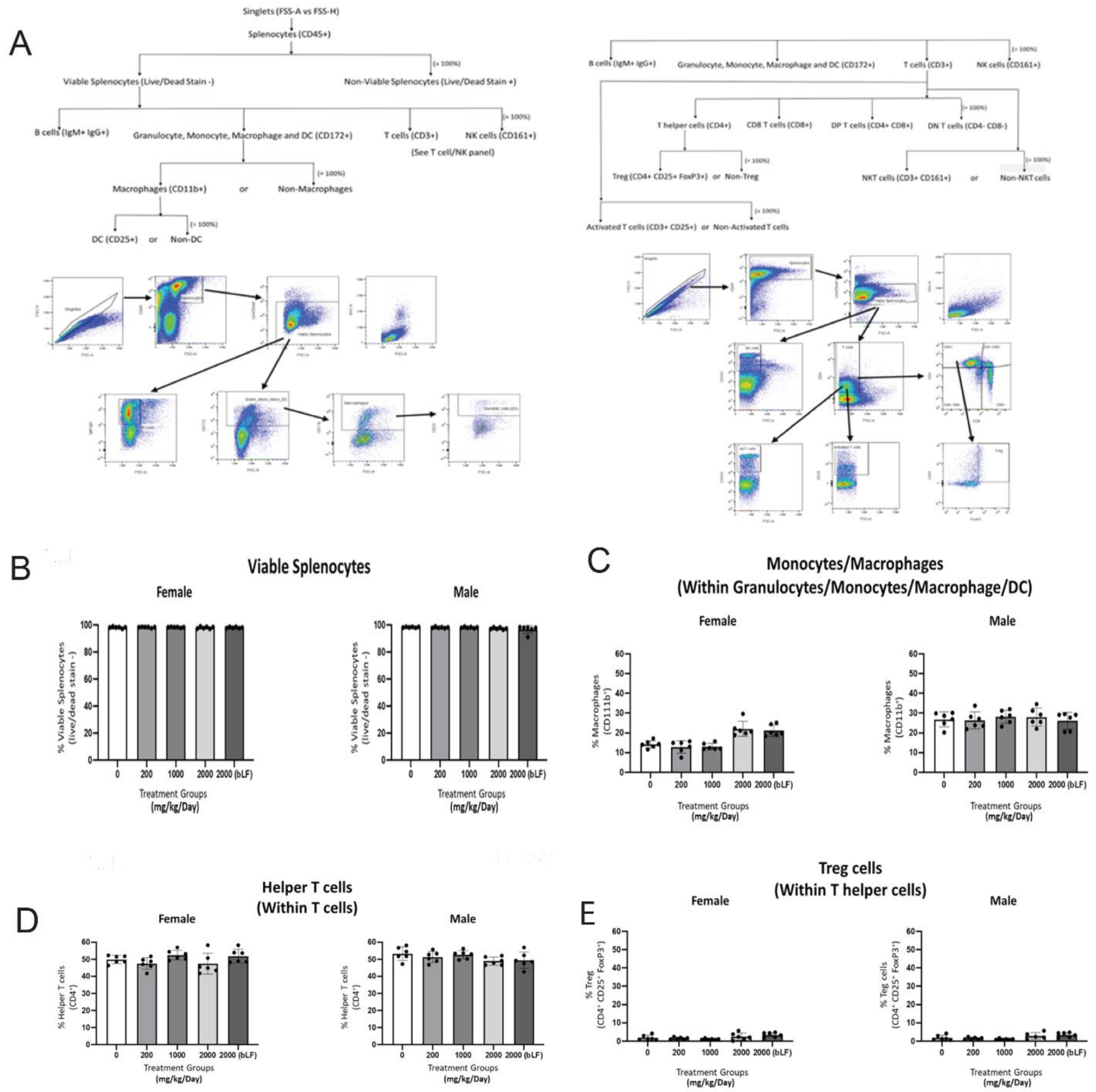


**Supplemental Figure 1.** Spleens were collected after the 14-day exposure period and dispersed into single-cell suspension for immunophenotyping analysis via flow cytometry. **A**. Overall gating strategy. **B**. Viable cells (within CD45+ splenocyte population). **C**. CD11b+ cells (within CD172a+ population). **D**. CD4+ helper T cells (within CD3+ T cell population). **E**. CD25+FoxP3+ regulatory T cells (within CD4+ T cells). N = 6 animals per group.
